# Topological Data Analysis of Human Brain Networks Through Order Statistics

**DOI:** 10.1101/2022.04.06.487253

**Authors:** Soumya Das, D. Vijay Anand, Moo K. Chung

## Abstract

Understanding the topological characteristics of the brain network across a population is central to understanding brain functions. The abstraction of human connectome as a graph has been pivotal in gaining insights on the topological features of the brain network. The development of group-level statistical inference procedures in brain graphs while accounting for the heterogeneity and randomness still remains a difficult task. In this study, we develop a robust statistical framework based on persistent homology using the *order statistics* for analyzing brain networks. The use of order statistics greatly simplifies the computation of the persistent barcodes. We validate the proposed methods using comprehensive simulation studies and subsequently apply to the resting-state functional magnetic resonance images. We conclude a statistically significant *topological* difference between the male and female brain networks.

**Author summary:** We fit a random graph model to the brain network and compute the expected persistent barcodes using *order statistics*. This novel approach significantly simplifies the computation of expected persistent barcodes, which otherwise requires complex theoretical constructs. Subsequently, the proposed statistical framework is used to discriminate if two groups of brain networks are topologically different. The method is applied in determining the *sexual dimorphism* in the shape of resting-state functional magnetic resonance images.

## Introduction

Modeling the human brain connectome as graphs has become the cornerstone of neuroscience, enabling an efficient abstraction of the brain regions and their interactions [1, 2]. Graphs offer the simplistic construct with only a set of nodes and edges to describe the connectivity of the brain network [3]. The generalizability of graph representation allows one to obtain quantitative measures across multiple spatio-temporal scales ranging from the node level up to the whole network level [4, 5]. To build the graph representation of brain networks, the whole brain is usually parcellated into hundreds of disjoint regions, which serves as nodes and the edges are associated with weights that indicate the strength of connection between the brain regions [6]. The graph theory based models provide reliable measures such as small-worldness, modularity, centrality and hubs [7–9]. However, these measures are often affected by the choice of arbitrary thresholds on the edge weights and thus make comparisons across networks difficult [10, 11]. To overcome this issue, the topological data analysis (TDA) has emerged to be a powerful method to systematically extract information from hierarchical layers of abstraction [12–15].

Persistent homology (PH), one of the TDA techniques, provides a coherent framework for obtaining topological invariants or features such as connected components and cycles at different spatial resolutions [16–19]. These topological invariants are often used to provide robust quantification measures to assess the topological similarity between networks [6]. Mostly the persistent barcodes are represented as persistent landscapes or diagrams and their distributions are used to compute a topological distance measure [20]. The PH based topological distances are found to consistently outperform traditional graph based metrics [21]. The main idea of using PH to brain networks is to generate a sequence of nested networks over every possible threshold through a graph filtration, which builds the hierarchical structure of the brain networks at multiple scales [10, 22–24].

In the graph filtration, a series of nested graphs containing topological information at different scales are produced. During the graph filtration, some topological features may live longer, whereas others *die* quickly. The filtration process tracks the birth, death and persistence of the topological features. The lifespans or *persistence* of these features are directly related to the topological properties of networks. The collection of intervals from births to deaths that defines persistences are called the barcodes which completely characterizes the topology of an underlying dataset [14]. The persistent diagram displays the paired births and deaths as scatter points [16, 20, 25, 26, 26–29]. The Betti curves, which counts the number of such features over filtrations, provide comprehensive visualizations of these intervals [6]. Thus, it is instructive to develop a statistical inference procedure using the persistent barcodes in order to compare across different groups and achieve meaningful inferences.

Since the real brain networks are often affected by heterogeneity and intrinsic randomness [30, 31], it is challenging to build a coherent statistical framework to transform these topological features as quantitative measures to compare across different brain networks by averaging or matching [32]. The brain networks are inherently noisy which makes it even harder to establish similarity across networks. Thus, there is a need to develop a statistical model that accounts for the randomness and provides consistent results across networks. The statistical models based on the distributions are expected to be more robust and less affected by the presence of outliers. To this end, we use the concept of random graph to analyze brain networks across a population.

Recently, there is a resurge of interest on using random graphs for network analysis. A graph whose features related to nodes and edges are determined in a random fashion is called a random graph. The theory of random graphs lies at the intersection between graph theory and probability theory. They are usually described using a probability distribution or a stochastic process that generates them [33, 34]. Analogous to the persistent barcodes, we also have their stochastic versions referred as the expected persistent barcodes for the random graphs. However, computing them requires complex theoretical constructs and they are generally approximated [35–37].

In this paper, we propose a more adaptable random graph model for brain networks. We consider a random complete graph, where all the nodes are connected with its edge weights randomly drawn from a continuous distribution. The consideration of a complete graph model simplifies building graph filtration straightforward [22, 32]. We then compute the expected 0D and 1D barcodes through the order statistics [38–43]. The use of order statistics in computing persistent homology features such as persistent barcodes and Betti numbers can drastically speed up the computation. Further, we propose the *expected topological loss* (ETL), which quantifies the 0D and 1D barcodes obtained through order statistics. We use the ETL as a test statistic to determine the topological similarity and dissimilarity between networks. The proposed random graph model and corresponding ETL methods are validated using extensive simulation studies with the ground truths. Subsequently, the method is applied to the resting-state functional magnetic resonance images (rs-fMRI) of the human brain.

## Materials and Methods

### Data

We considered a resting-state fMRI dataset collected as part of the Human Connectome Project (HCP) [44, 45]. The dataset consisted of the subset of fMRI scans of 400 subjects (168 males and 232 females) over approximately 14.5 minutes using a gradient-echoplanar imaging sequence with 1200 time points [24, 32]. Informed consent was obtained from all participants by the Washington University in St. Louis institutional review board [46]. The ethics approval for using the HCP data was obtained from the local ethics committee of University of Wisconsin-Madison.

The human brain can be viewed as a weighted network with its neurons as nodes. However, considering a high number of neurons (~ 10^12^) in a human brain, the traditional brain imaging studies parcellate the brain into a manageable number of mutually exclusive regions [47–49]. These regions are then considered as nodes while the strength of connectivity between these regions are edges. For the considered dataset, the Automated Anatomical Labeling (AAL) template was employed to parcellate the brain volume into 116 non-overlapping anatomical regions [50] and the fMRI across voxels within each brain parcellation were averaged. This resulted 116 average fMRI time series with 1200 time points for each subject. Further, we removed fMRI volumes with significant head movements [51] because such movements are shown to produce spatial artifacts in functional connectivity [52–54].

### Simplicial complex

A *simplex* is a generalization of the notion of a triangle or tetrahedron to arbitrary dimensions. A 0-simplex is a point, a 1-simplex is a line segment, and a 2-simplex is a triangle. In general, a *k*-simplex *S_k_* is a convex hull of *k* + 1 affinely independent points 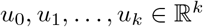:

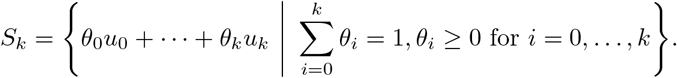

Whereas, a simplicial complex 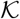 is a set of simplices that satisfies the following two conditions. (1) Every face of a simplex from 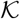 is also in 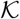. (2) The non-empty intersection of any two simplices *S*_1_, 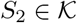 is a shared face [55]. We call a simplicial complex consisting of up to *k*-simplices a *k*-skeleton. Since graphs are a collection of nodes (0-simplices) and edges (1-simplices), they are 1-skeleton simplicial complexes.

### Graph and graph filtration

Consider a weighted graph *G*(*p*, ***w***), where *p* is the number of nodes and ***w*** = (*w*_1_,…, *w_q_*)^⊤^ is a *q*-dimensional vector of edge weights with *q* = (*p*^2^ – *p*)/2. The binary graph *G_ϵ_*(*p*, ***w**_ϵ_*) of *G*(*p*, ***w***) has binary edge weight *w*_*ϵ*,*i*_:

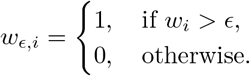

The binary network *G_ϵ_*(*p*, ***w**_ϵ_*) is a 1-skeleton. In 1-skeleton, 0-dimensional (0D) holes are *connected components* while the 1-dimensional (1D) holes are *cycles*. There is no higher dimensional homology beyond dimensions 0 and 1 in 1-skeleton. The number of connected components and the number of independent cycles in a graph are referred to as the 0th Betti number (*β*_0_) and 1st Betti number (*β*_1_), respectively. For 1-skeletons, there is an efficient 1D filtration method called the *graph filtration*, which filters at the edge weights from –∞ to ∞ in a sequentially increasing manner [6, 32]. The graph filtration of *G* is defined as a collection of nested binary networks

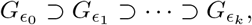

where *ϵ*_0_ < *ϵ*_1_ < ··· < *ϵ_k_* are filtration values. We consider the edge weights as the filtration values to make the graph filtration unique.

### Birth and death decomposition

When we increase the filtration value *ϵ*, either one new connected component appears or one cycle disappears [6]. Once a connected component is born, it never dies implying an infinite death value. On the other hand, all the cycles are considered to be born at –∞. Thus, the number of connected components (or cycles) is non-decreasing (or non-increasing) as *ϵ* increases. Subsequently, the 0D barcode is given by a set of increasing birth values:

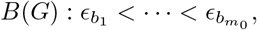

and the 1D barcode is given by a set of increasing death values:

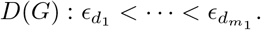

The 0D and 1D barcodes are often visualized using persistent diagrams [16, 25–27] and Betti curves. The Betti curves plot the Betti numbers with respect to the filtration values. Since the Betti numbers are monotonic, the Betti curve is a step function with a one-unit jump (or drop) at every birth (or death) values. We can show that the total number of finite birth values of connected components and the total number of death values of cycles are

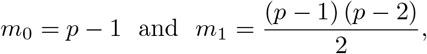

respectively [32]. The number of connected components (*β*_0_) and cycles (*β*_1_) in a complete graph *G*_0_ are equal to 1 and *m*_1_, respectively. We note that every edge weight must be in either 0D barcode or 1D barcode as summarized in the following theorem [32].

#### Theorem 1 (Birth-death decomposition)

*The set of* 0*D birth values B(G) and* 1*D death values D*(*G*) *partition the edge weight vector **w** such that B*(*G*) ∪ *D*(*G*) = ***w** and B*(*G*) ⋂ *D*(*G*) = *ϕ. The cardinalities of B*(*G*) *and D*(*G*) *are p* – 1 *and* (*p* – 1) (*p* – *2*)/*2*, *respectively*.

### Wasserstein distance on barcodes

Since the barcodes completely characterize the topology of a network [14], the topological similarity between two such networks can be quantified using a distance metric between the corresponding 0D or 1D barcodes [56]. Often used metric is the Wasserstein distance [23, 57–59]. Let *G*_1_ (*p*, ***u***) and *G*_2_ (*p*, ***v***) be two networks and the corresponding barcodes (or persistent diagrams) be *P*_1_ and *P*_2_. Then, the 2-*Wasserstein distance* on barcodes is given by

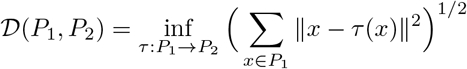

over every possible bijection *τ* between *P*_1_ and *P*_2_ [23, 60, 61]. For graph filtrations, barcodes are 1D scatter points. Therefore, the bijection *τ* can be simplified to the 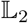 norm between the sorted birth values of connected components or the sorted death values of cycles [23].

#### Theorem 2

*Let G*_1_ *and G*_2_ *be two networks with p nodes and*

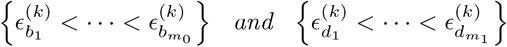

*be the birth and death sets of the network G_k_*, *k* = 1, 2. *Then*, *the* 2-*Wasserstein distance between the* 0D *barcodes for graph filtration is given by*

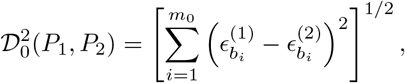

*and the* 2-*Wasserstein distance between the* 1D *barcodes is*

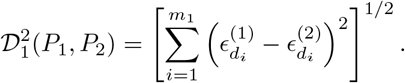

### Expected persistent barcodes of random graph

We consider a random graph 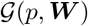, where its edge weights are drawn independent and identically from a distribution function *F_W_*. Here, *p* is the number of nodes and ***W*** = (*W*_1_,…, *W_q_*)^⊤^ is a q dimensional vector of random weights with *q* = (*p*^2^ – *p*)/2. Fig 1 displays weighted brain networks randomly drawn from Beta distributions. The considered graph is complete and its edge weights are drawn randomly from a continuous distribution. To be mathematically precise, the considered random graph is *almost surely* complete. Since we assume the edge weights to be drawn from a continuous distribution, the probability of a certain edge weight being zero is *nil*.

If we apply a graph filtration on the random weighted graph 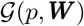, we have a set of random birth values of connected components (or random 0D barcode) and a set of random death values of cycles (or random 1D barcode). Since the notions of random birth and death values are abstract, it is important to turn them into deterministic topological descriptors. As often, one of the simplest way to turn a random object into a deterministic summary is to consider its average behavior. To that end, we study the expected birth and death values (or expected persistent barcodes) as follows.

#### Expected birth and death values

Let 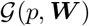 be a random graph and its sorted *random* edge weights be

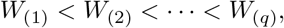

where the subscript (*i*) indicates the ith smallest *random* edge weight. Let the *random* birth and death values of the connected components and cycles be

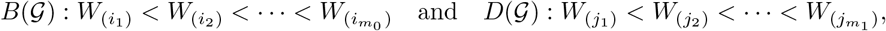

where *m*_0_ = *p* – 1 and *m*_1_ = (*p* – 1)(*p* – 2)/2. Then, the expected birth and death values are given by

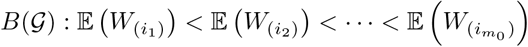

and

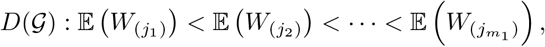

where 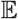 indicates the standard *expectation operator* on a *random* weight.

In order to compute the expected birth and death values, we provide an explicit expression for 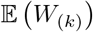, for *k* = 1,…, *q*, through Theorem 3 below.

##### Theorem 3

*Let the edge weights **W*** = {*W*_1_, *W*_2_,…, *W_q_*} *of a random graph* 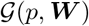 *be drawn from a distribution with cumulative distribution function (cdf) F_W_ and probability density function (pdf) f_W_. Then, the expectation of the kth order statistic can be approximated by*

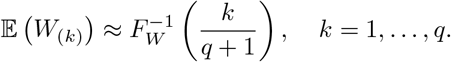

**Proof** Since the edge weights *W*_1_, *W*_2_,…, *W_q_* are drawn from a distribution with a cdf *F_W_* and a pdf *f_W_*, the pdf of the *k*th order statistic *W*_(*k*)_ is given by

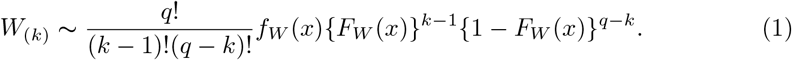

*W*_(*k*)_ does not follow a well-known distribution and, therefore, the computation of its mean and variance is difficult. However, [62] showed that the *r*th sample quantile of {*W*_1_, *W*_2_,…, *W_q_*} is asymptotically normally distributed:

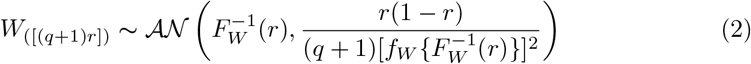

for large *q*, where 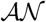 stands for *asymptotic normal* distribution. Thus, the approximate mean and variance of *W*_(*k*)_ can be found from (2) by letting *r* = *k*/(*q* +1):

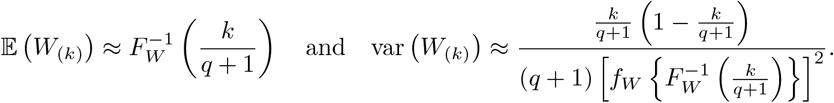

(Here we also compute the variance as we shall use that later while computing confidence intervals.) Hence, the proof is complete.

Now, we use Theorem 3 and provide expressions for the expected birth and death values in Theorem 4 below.

##### Theorem 4

*Let* 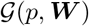 *be a random graph, where its edge weights are drawn from a cdf F_W_. Then, the expected birth values of the connected components of* 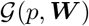 *are given by*

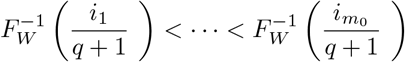

*and the expected death values of the cycles of* 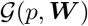 *are given by*

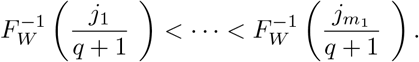

**Proof** The proof is trivial as the expected birth and death values are defined as

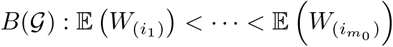

and

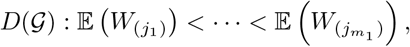

and Theorem 3 suggests

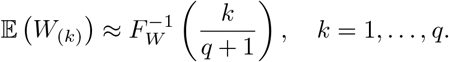

Below we present a corollary, where we consider a particular scenario of Theorem 4. We show that, if the edge weights follow a uniform distribution, then the expected birth and death values have a more simplified and an exact form.

#### Corollary

Let 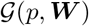 be a random graph with its edge weights drawn from a Uniform(0, 1) distribution. Then, the expected birth values of the connected components of 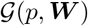 are given by

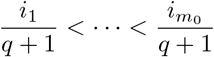

and the expected death values of the cycles of 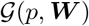 are given by

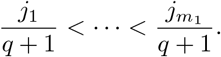

**Proof** If the weight distribution is Uniform(0, 1), then the pdf (1) of the kth order statistic simplifies to

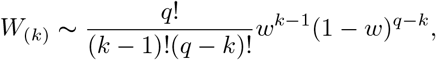

which turns out to be the pdf of the well-known Beta distribution with parameters *k* and *q* + 1 – *k*. Since the mean of a Beta(*k*, *q* + 1 – *k*) distribution has an exact form of *k*/(*q* + 1), we have

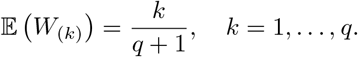

This completes the proof.

Once the expected birth and death values are computed, we use them to plot Betti curves as illustrated in Fig 3. For this purpose, we consider two random graphs with *p* = 150 nodes and their edge weights drawn from Beta(2, 2) and Beta(2, 3) distributions. We observe that a slight change in distribution significantly affects the topology of a network.

**Fig 1.**
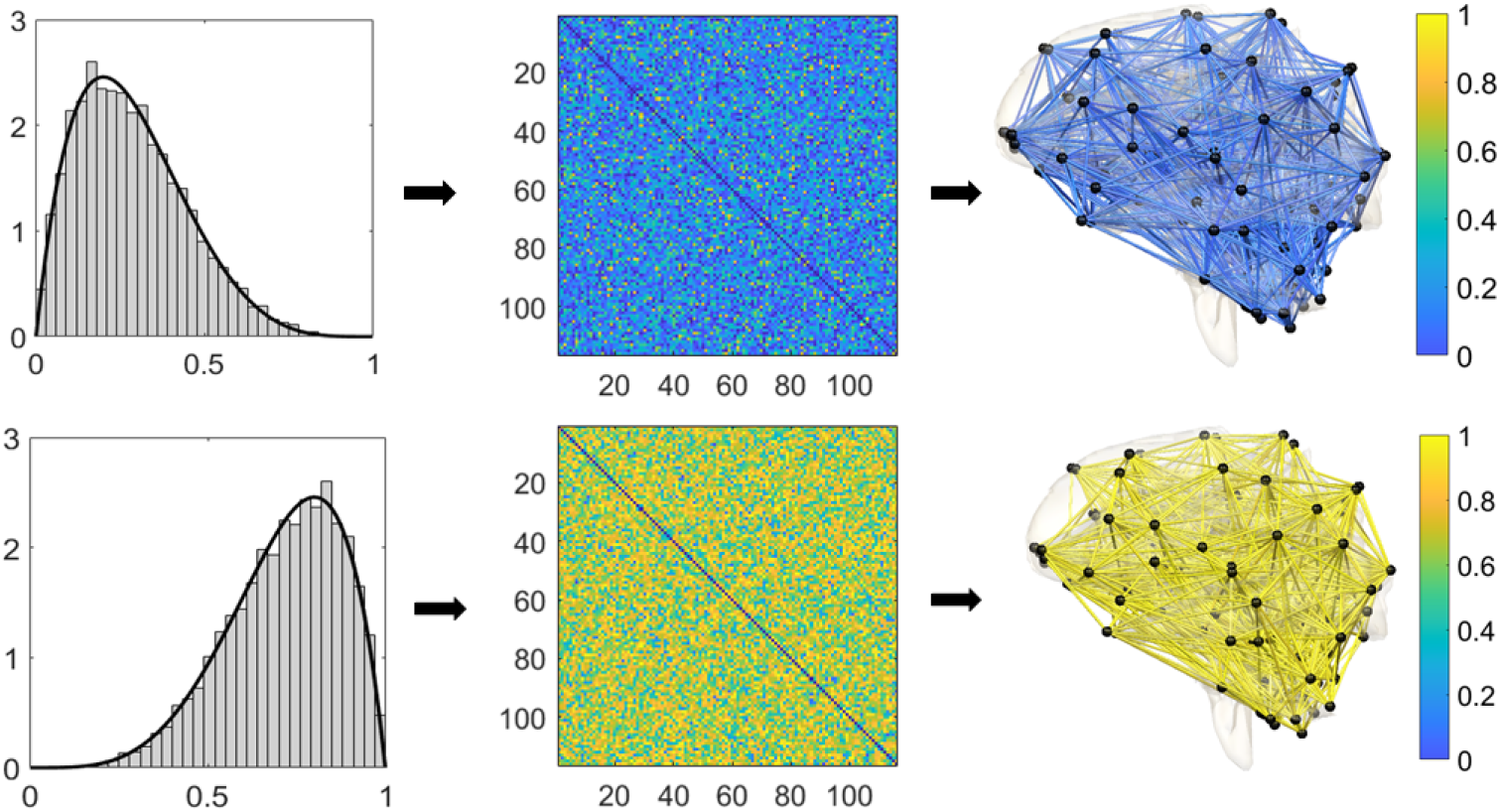
Visualization of simulated brain networks with 116 nodes. Left: The empirical density functions of simulated edge weights from Beta(2, 5) (top) and Beta(5, 2) (bottom) distributions. Middle: The 116 × 116 correlation matrices constructed using the simulated edge weights. Right: Human brain networks with the simulated edge weights. Since correlation networks are too dense for visualization, we only displayed edges with values below 0.1 and above 0.9.

**Fig 2.**
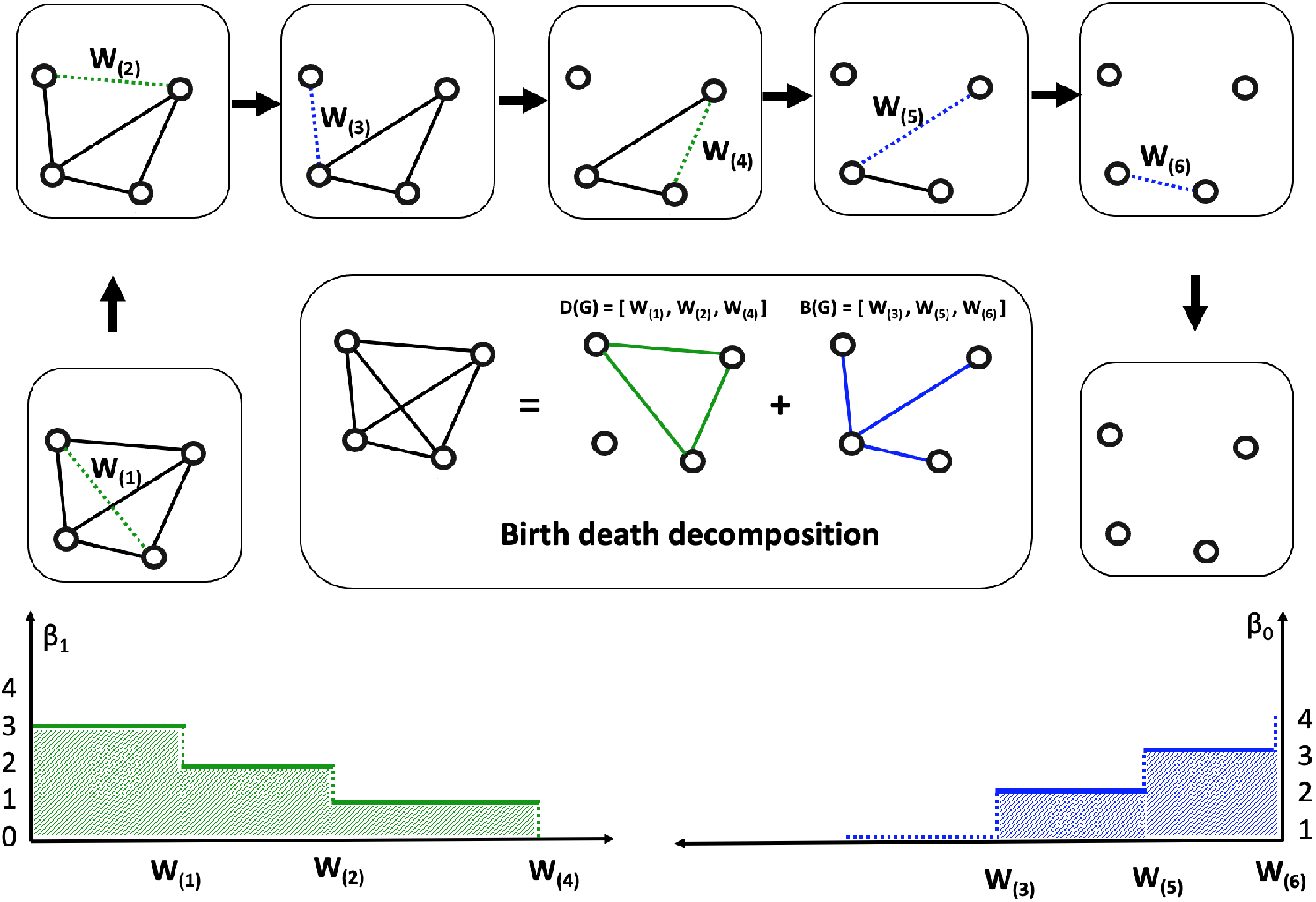
Schematic of graph filtration and persistent barcodes computation. We consider a random weighted graph with *p* = 4 nodes, where the number of edges is *q* = *p*(*p* – 1)/2 = 6. The random edge weights are {*W*_1_, *W*_2_,…, *W*_6_}. We order them using the order statistic as *W*_(1)_ < *W*_(2)_ < ··· < *W*_(6)_. We remove each edge of the random graph one at a time in the graph filtration and construct the random birth and death sets of the connected components and cycles, respectively. The Betti-0 (lower right) and Betti-1 (lower left) curves are drawn using the birth and death sets. The blue and green shaded areas represent the areas under Betti-0 and Betti-1 curves. Later, we will consider the area under Betti-0 curve to quantify the curve and construct a test statistic to discriminate between two groups of networks.

**Fig 3.**
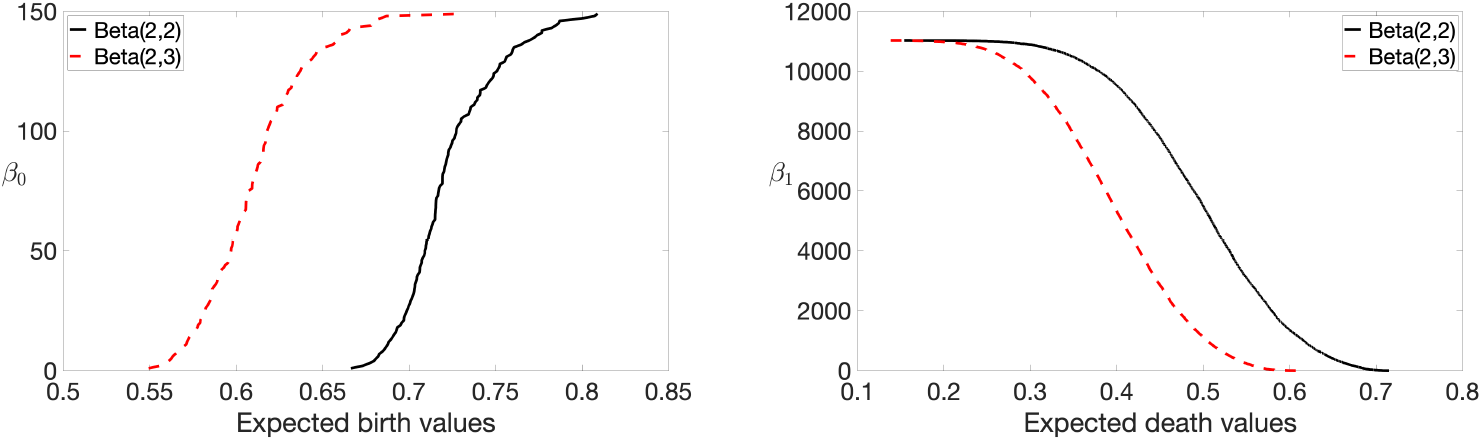
Plots of Betti-0 (left) and Betti-1 (right) curves of random networks with edges drawn from Beta(2, 2) (in dotted red line) and Beta(2, 3) (in solid black line) distributions. We observe that a slight change in distribution significantly affects the topology of a network.

### Estimation of birth and death values and their confidence bands

Given a set of *n* samples from a random graph 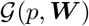, we present a statistical methodology to estimate expected birth and death values, and the corresponding confidence bands. The methodology is later validated using a simulation study. Let the random weights of the *n* sampled graphs be ***w***_1_, ***w***_2_,…, ***w**_n_*, where ***w**_i_* = (*w*_*i*1_, *w*_*i*2_,…, *w_iq_*)^τ^ and *w_ij_*~*F_W_* for all *i* = 1,…, *n* and *j* = 1,…, *q*.

From the previous section, we know that *W*_(*k*)_ follows a asymptotic Gaussian distribution with its mean and variance being

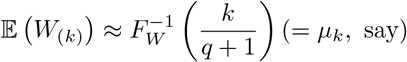

and

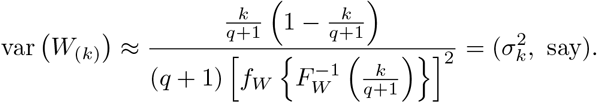

Thus, computing the mean and variance requires estimating the pdf *f_W_* and the inverse cdf 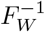. To estimate *f_W_*, we average the *n* graphs (with respect to their weights) and consider the Gaussian kernel density estimate (KDE) of the averaged weights. Let the average weight vector be 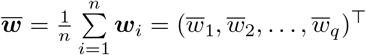. Then, the KDE of the pdf *f_W_* is given by [63]

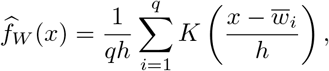

where *K* is a non-negative kernel and *h* > 0 is a smoothing parameter called *bandwidth*. For the current purpose, we assume the kernel to be Gaussian, i.e., *K*(·) = *ϕ*(·).

To estimate 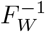, we first find the empirical cdf of *F_W_* based on the averaged weight vector 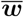:

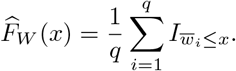

Here, *I_A_* is an indicator function that takes value 1 if the event *A* is true and 0 otherwise. Then, we calculate the inverse cumulative distribution of 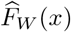:

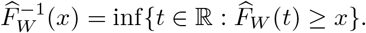

Once *f_W_* and 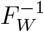 are estimated, we plug-in the corresponding estimates in *μ_k_* and 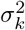 to have 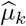 and 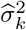, for *k* = 1,…, *q*. Finally, we calculate the *α*% confidence intervals as

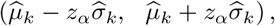

where *z_α_* is such that

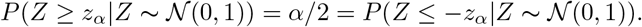

In particular, if *α* = 95, we have *z_α_* = 1.96.

### Inference on expected birth and death values

Since a graph can be topologically characterized by 0D and 1D barcodes, the topological similarity and dissimilarity between two graphs can be measured using the differences of such barcodes. To quantify these differences, we propose *expected topological loss (ETL)* as follows.

Let 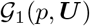 and 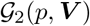 be two random graphs, where the random weights ***U*** = {*U*_1_,…, *U_q_*} and **V** = {*V*_1_,…, *V_q_*} are drawn from distribution functions *F_U_* and *F_V_*, respectively. Further, let the expected birth and death values of 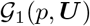 be

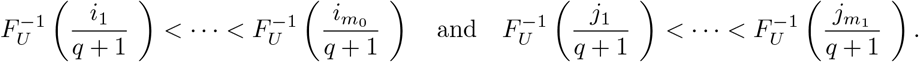

Similarly, let the expected birth and death values of 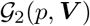 be

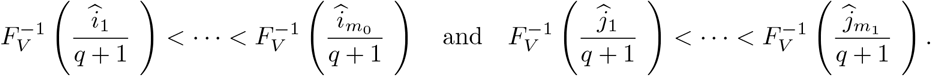

Then, the ETL is given by

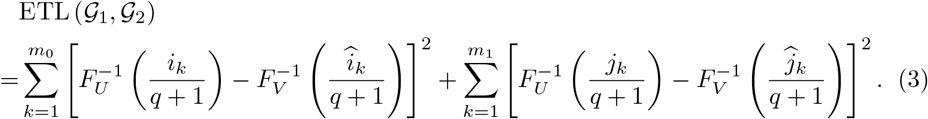

In most real life scenarios, the distribution functions *F_U_* and *F_V_* are unknown. In such scenarios, we plug-in the corresponding empirical distribution function estimates, 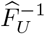 and 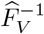, in (3).

### Application of ETL in discriminating two groups

The ETL can be used to topologically discriminate between two groups of brain networks. Let **Ω** = {Ω_1_,…, Ω_*m*_} and **Ψ** = {Ψ_1_,…, Ψ_*n*_} be two sets consisting of *m* and *n* complete networks each comprising *p* number of nodes. Further, let the empirical distribution functions of the edge weights of the graphs in group **Ω** be

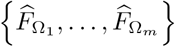

and the expected birth and death values for Ω_*i*_, *i* = 1,…, *m*, be

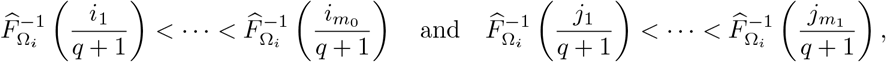

which, for simplicity, we denote by

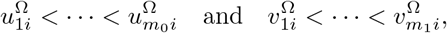

respectively. Thus, the average (standard mean) of expected birth and death values for the group **Ω** are given by

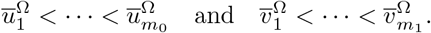

Similarly, for the second group **Ψ**, let the average of expected birth and death values be

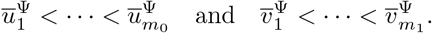

Now, we consider a statistic based on ETL to discriminate between groups:

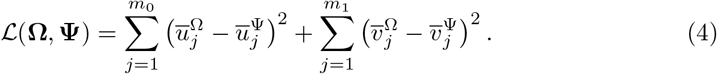

A “large” 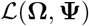 shall indicate a significant topological difference between the two groups whereas a “small” value shall suggest that there is no significant topological group differences. Considering the probability distribution of the test statistic 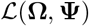 is unknown, we use the permutation test [64–67]. In this study, we use 100000 permutations. The *p*-values are computed after 10 such independent simulations.

A similar widely-used statistic is the maximum gap statistics. On a similar line to 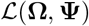, the statistic is given by:

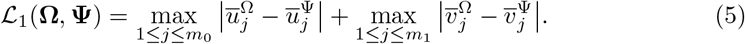

We will use 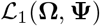 to compare with the ETL statistic 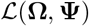 in the simulation study section.

### Area under Betti-0 curve in discriminating groups

The characteristics of *β*_0_ can be quantified using the area under the Betti-0 curve (AUC) [68]. Using the notations of the previous subsection, the AUC for Ω_*i*_ of the group **Ω** and for Ψ_*j*_ of the group **Ψ** are given by

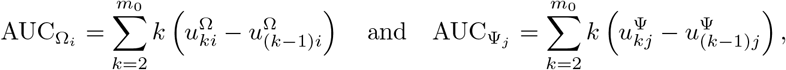

for *i* = 1,…, *m* and *j* = 1,…, *n*. We compute the AUC by summing up the areas of rectangular blocks formed by the expected persistent barcodes. For example, if we consider Fig 2, the area under the Betti-0 curve is

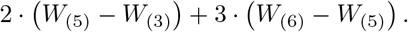

In this scenario, we only have two terms because *m*_0_ = *p* – 1 = 3.

To determine if AUC is significantly different between the groups **Ω** and **Ψ**, we consider the Wilcoxon rank sum test [69]. The Wilcoxon rank sum test is a nonparametric test of the null hypothesis that, for randomly selected values *X* and *Y* from two populations, the probability of *X* being greater than *Y* is equal to the probability of *Y* being greater than *X*. This is unlike the previous situation, where we considered a ETL or a max-gap statistic. In those scenarios, since we are considering the distance between increasing births (or deaths) of two networks (with equal number of nodes), the consideration of 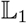 or 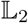 norm in the statistic is meaningful. Whereas, in the current scenario, we have two samples of AUC of networks of size *m* and *n*. Therefore, a distance statistic cannot be employed.

The Wilcoxon rank sum test places ranks to the aggregated sample (combined first and second sample) and considers the sum of ranks for both the samples. For *p* > 4, the number of cycles increases in power scale compared to the number of connected components. Since the number of cycles is generally quite higher than the number of connected components, the area under *β*_1_ curve dominates the total area. Also, while comparing two samples coming from two different distributions, the areas under *β*_1_ curves differ significantly. Thus, the Wilcoxon rank sum test puts all the lower ranks (say, 1 to *m*) to one of the samples and the remaining ranks to the other (say, *m* + 1 to *m* + *n*). This makes the statistic constant even if we vary the distributions. This implies the sum of ranks are same as long as we are considering two significantly different distributions. Therefore, the results are robust (the test can identify difference as long as there is a difference in distributions) but the *p*-values are always same for different set of distributions (because the sum of ranks are same). For example, the *p*-value is always exactly 0.0022 if we test the difference between two groups with 6 networks each, where the edge weights are drawn from “Beta(1,1) & Beta(5,2)” or “Beta(1,5) & Beta(5,2)” or “Beta(1,1) & Beta(1,5).” To make the test more unpredictable, we are only considering the area under *β*_0_ curve. However, one may always incorporate the area under *β*_1_ curve as it is more robust and the results are stable.

### Simulation study

#### Validation of birth and death value estimates

We validate the method to estimate expected birth and death values. We consider a (non-stochastic) graph *G*(*p*, ***w***) with given edge weights and calculate its birth and death values using the standard method [32]. On the other hand, we simulate *n* vectors of *q*-variate Gaussian noises and add them to the edge weights ***w*** of *G*(*p*, ***w***) to have a set of *n* graphs

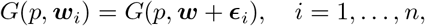

where 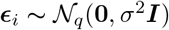. We consider the hence produced set of graphs

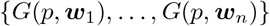

as realizations from a random graph 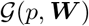 and apply the proposed method to calculate the expected birth and death values along with their corresponding confidence bands. Then, we compare them with the initially calculated birth and death values of *G*(*p*, ***w***). A schematic of the validation procedure is presented in Fig 4.

**Fig. 4.**
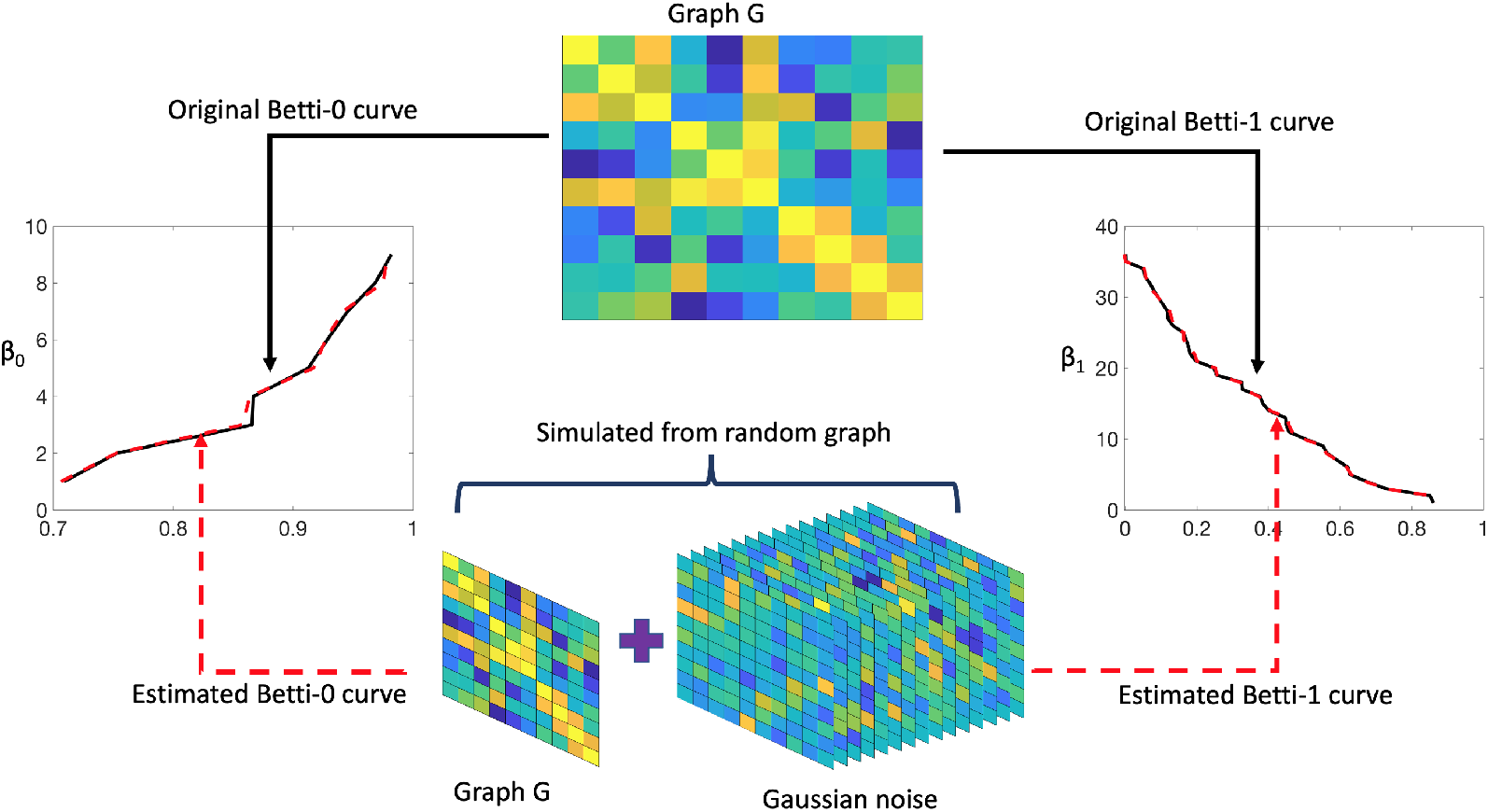
Schematic to validate the proposed method to compute expected birth and death values and, hence, the Betti curves. The graph *G* is a non-stochastic graph, which we used to calculate the Betti curves (solid black line) in the standard way. On the other hand, we added 15 Gaussian noise-graphs to the main graph *G* to have a set of 15 graphs. These 15 graphs are assumed to be sampled from a random weighted graph. Then, we apply the proposed method on this set of sampled graphs and estimate the expected birth and death values and, hence, the Betti curves (dotted red line).

To generate the graph *G*(*p*, ***w***), we considered *p* =10 number of nodes. That is, the weight vector ***w*** is of dimension *q* = *p*(*p* – 1)/2 = 45. We drew the *q* variate random weights ***w*** from the Uniform(0, 1) distribution. We simulated *n* = 15 noise vectors from 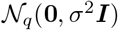 of dimension *q* with *σ* = 0.02. The original and expected birth (left panel) and death (right panel) values are plotted in Fig 5. The black line represents the original birth (or death) values, the dashed red line indicates the estimated birth (or death) values, and the dashed blue lines indicate the corresponding 95% upper and lower confidence bands. We observe that the dashed red lines almost overlap the black lines of the original birth and death values. In addition, the original birth (or death) values almost always lie within the confidence bands supporting the reliability of the proposed methodology.

**Fig. 5.**
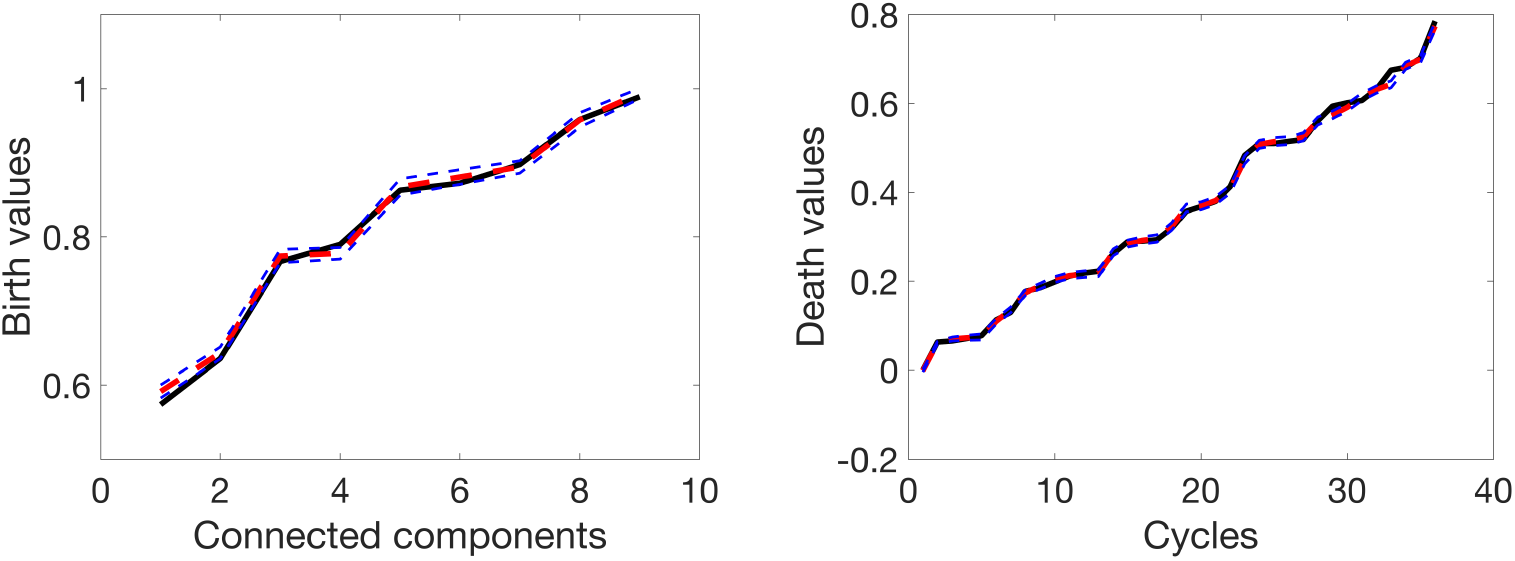
Plots of the original and expected birth (left) and death (right) values. The black line represents the original birth (or death) values, the dashed red line indicates the expected birth (or death) values, and the dashed blue lines indicate the corresponding 95% confidence intervals.

#### Analyzing topological similarity between two groups of brain networks

##### Using expected topological loss

We provide a toy example to illustrate whether the topological similarity (or dissimilarity) of two groups of networks, drawn from two different distributions (or the same distribution), can be identified using the ETL statistic (4). To that end, we consider the Uniform(0, 1) and Beta(*a*, *b*) distributions. Both the distributions are defined on the domain (0, 1). The shape parameters *a* and *b* of the beta distribution allow it to take a variety of shapes including the shape of a uniform pdf when *a* = *b* = 1. We consider two different shapes, other than the uniform one, for validation. The top panel of Fig 6 demonstrates these shapes of the beta density functions.

**Fig. 6.**
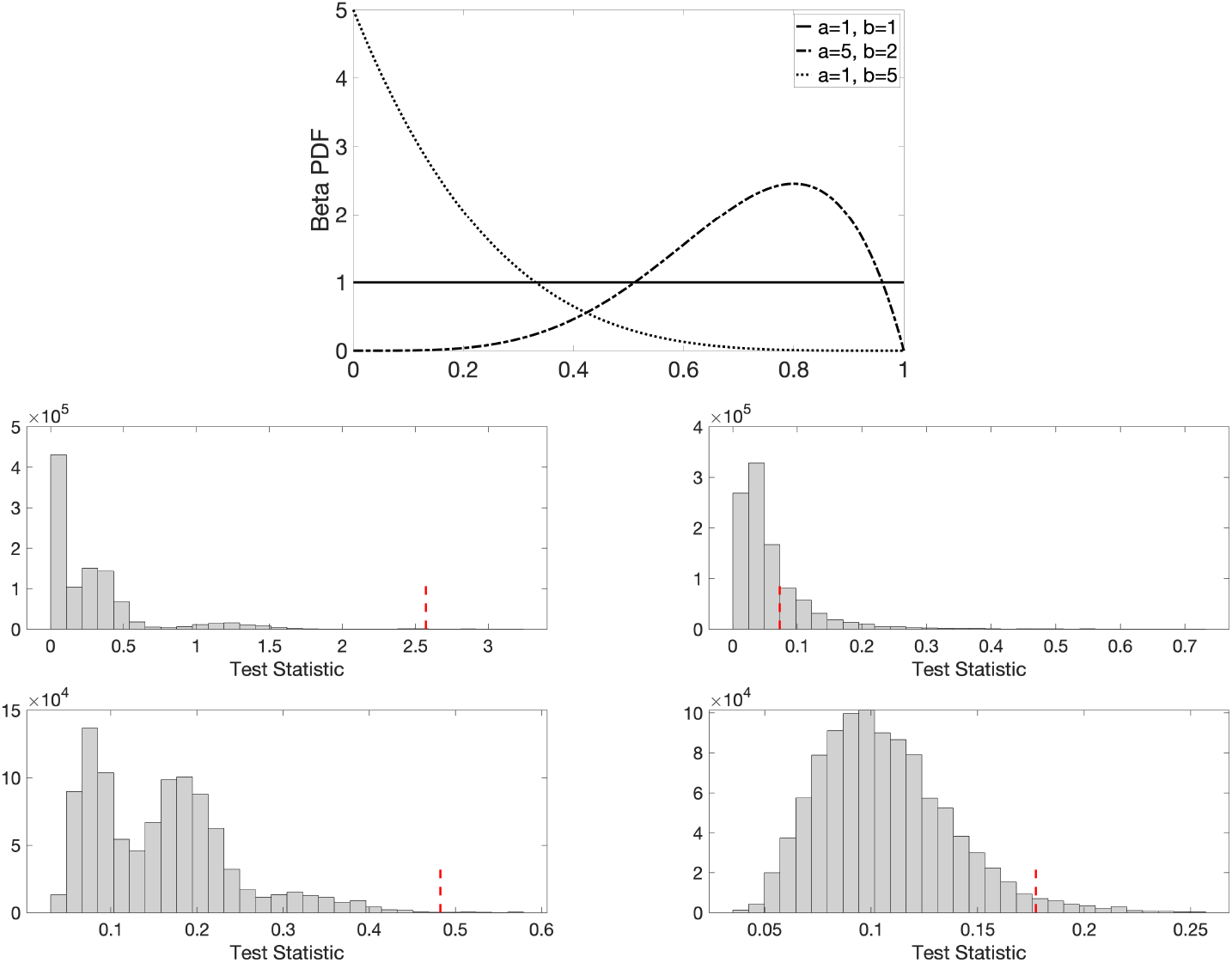
Top panel: Probability density functions of Beta(1, 1) or Uniform(0, 1) (in solid line), Beta(5, 2) (in dash-dotted line), and Beta(1, 5) (in dotted line). We sample the edge weights of random graphs from these three different distributions for validation purpose. Middle and bottom panel: Histogram plots of the ETL (middle) and maximum gap (bottom) test statistics and the corresponding observed test statistics (in dotted red lines) for the scenarios: Beta(1, 1) vs. Beta(5, 2) (left) and Beta(1, 1) vs. Beta(1, 1) (right) with 6 networks in each group.

For both the groups, we simulated *n* = 6, 8, 10, and 12 networks. Each network was constructed using *p* =10 nodes. That is, we had *q* = *p*(*p* – 1)/2 = 45 number of edges, *m*_0_ = *p* – 1 = 9, and m_1_ = (*p* – 1)(*p* – 2)/2 = 36. For the permutation test, we considered 100000 permutations and repeated that 10 times to compute the average *p*-values. Table 1 tabulates the *p*-values for different pairs of considered distributions. In all the scenarios, networks drawn from the same distribution produced large *p*-values and networks drawn from different distributions had *p*-values smaller than 0.01. Therefore, we conclude that the proposed ETL statistic, based on expected birth and death values, can discriminate networks drawn from different distributions at 99% confidence level. The middle panel of Fig 6 plots the histograms of the ETL test statistic and the corresponding observed test statistics (in dotted red) for two specific scenarios: (i) Beta(1, 1) vs. Beta(5, 2) (left) and (ii) Beta(1, 1) vs. Beta(1, 1) (right) with 12 networks in each group.

**Table 1.**
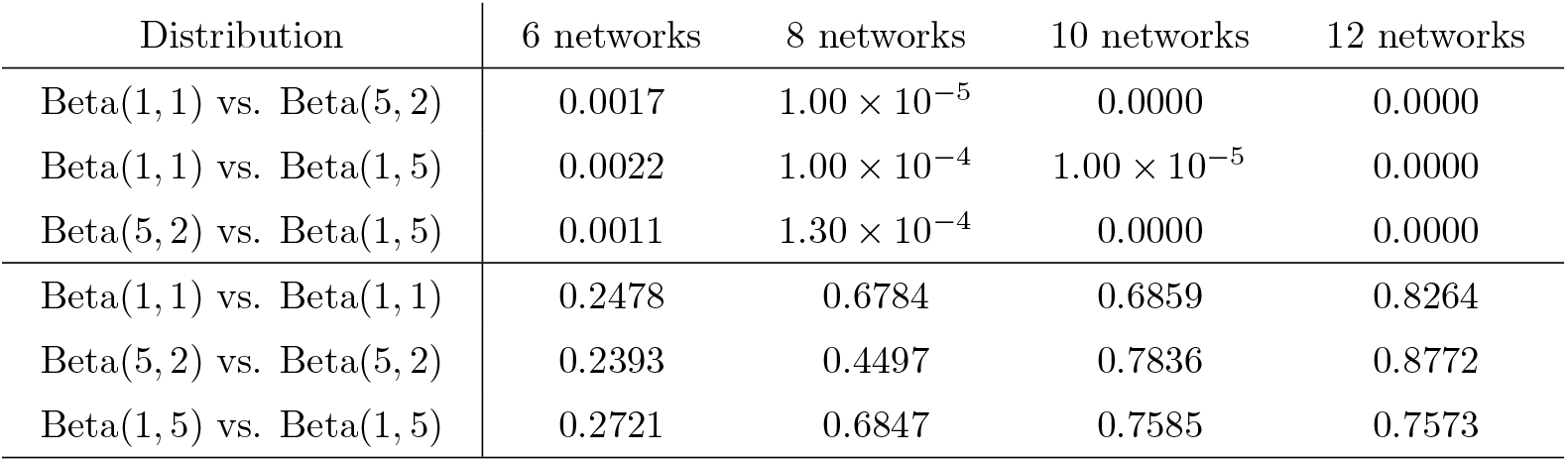
The average *p*-values obtained using the **ETL statistic** for various pairs of distributions considered for drawing edge weights of networks. Here, the columns 6 networks, 8 networks, 10 networks, and 12 networks indicate the number of networks that we considered for both the groups. The *p*-values smaller than 0.01 indicate that our method can identify network differences at a 99% confidence level.

| Distribution | 6 networks | 8 networks | 10 networks | 12 networks |
| --- | --- | --- | --- | --- |
| Beta(1, 1) vs. Beta(5, 2) | 0.0017 | $1.00 \times 10^{-5}$ | 0.0000 | 0.0000 |
| Beta(1, 1) vs. Beta(1, 5) | 0.0022 | $1.00 \times 10^{-4}$ | $1.00 \times 10^{-5}$ | 0.0000 |
| Beta(5, 2) vs. Beta(1, 5) | 0.0011 | $1.30 \times 10^{-4}$ | 0.0000 | 0.0000 |
| Beta(1, 1) vs. Beta(1, 1) | 0.2478 | 0.6784 | 0.6859 | 0.8264 |
| Beta(5, 2) vs. Beta(5, 2) | 0.2393 | 0.4497 | 0.7836 | 0.8772 |
| Beta(1, 5) vs. Beta(1, 5) | 0.2721 | 0.6847 | 0.7585 | 0.7573 |

##### Comparison of expected topological loss with baseline approaches

We compared the proposed ETL with several other widely-used baseline topological distances such as bottleneck, Gromov-Hausdorff (GH), and Kolmogorov-Smirnov (KS) distances [21, 70, 71]. We also compared the results with the maximum gap statistic defined earlier in (5). In all the scenarios, we considered two groups of networks each of size *n* = 6. The remaining simulation setting is similar to the above. The corresponding *p*-values are presented in Table 2. From the table, we observe that the ETL performs well in most scenarios. In particular, we note that the KS based methods do not perform well whereas the maximum gap based method is quite competitive. Further, for testing no network differences, all the distances perform well.

**Table 2.** The average *p*-values obtained using bottleneck, GH, KS, maximum gap, and ETL based statistics for various pairs of distributions considered for drawing edge weights of networks. There were 6 networks in each group for all the scenarios. The *p*-values smaller than 0.01 indicate that the corresponding method can identify network differences at a 99% confidence level.

| Distribution | Bottleneck | GH | KS($\beta_0$) | KS($\beta_1$) | Maximum gap | ETL |
| --- | --- | --- | --- | --- | --- | --- |
| Beta(1, 1) vs. Beta(5, 2) | 0.0035 | 0.0028 | 0.4667 | 0.3438 | 0.0022 | 0.0014 |
| Beta(1, 1) vs. Beta(1, 5) | 0.0190 | 0.0692 | 0.3804 | 0.6406 | 0.0016 | 0.0022 |
| Beta(5, 2) vs. Beta(1, 5) | 0.0026 | 0.3345 | 0.2885 | 0.5177 | 0.0021 | 0.0011 |
| Beta(1, 1) vs. Beta(1, 1) | 0.2255 | 0.4385 | 0.2591 | 0.6893 | 0.1013 | 0.3136 |
| Beta(5, 2) vs. Beta(5, 2) | 0.3046 | 0.2346 | 0.1991 | 0.6035 | 0.1446 | 0.2393 |
| Beta(1, 5) vs. Beta(1, 5) | 0.3351 | 0.5392 | 0.1217 | 0.4172 | 0.1058 | 0.2721 |

Since the maximum gap based method exhibits a competitive performance with the ETL based method, we plot the histograms of the maximum gap statistics obtained over different permutations and the corresponding observed test statistics (in dotted red) for two specific scenarios: (i) Beta(1, 1) vs. Beta(5, 2) (left) and (ii) Beta(1, 1) vs. Beta(1, 1) (right) with 6 networks in each group; see the bottom panel of Fig 6. Although both the methods (ETL and maximum-gap) perform well, the ETL generally produces better results (i.e., its *p*-value is closer to 0 when there is a network difference and closer to 1 when there is no network difference).

##### Using area under Betti-0 curve

Finally, we conducted a simulation study for the method based on the area under *β*_0_ curve. The considered simulation layout was the same as before. The obtained *p*-values are tabulated in Table 3. Networks drawn from the same distribution produced large *p*-values and networks drawn from different distributions had *p*-values smaller than 0.01. Therefore, similar to the ETL, this approach can discriminate networks drawn from different distributions at a 99% confidence level.

**Table 3.**
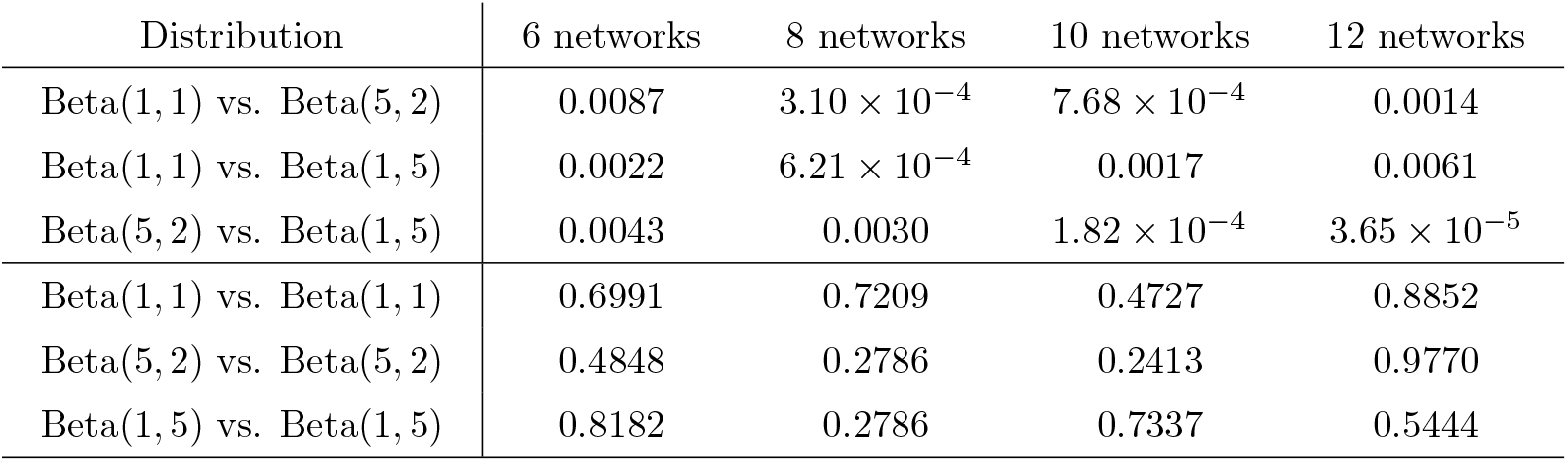
The average *p*-values obtained using Wilcoxon rank sum test on the areas under *β*_0_ curves for various pairs of distributions considered for drawing edge weights of networks. Here, the columns 6 networks, 8 networks, 10 networks, and 12 networks indicate the number of networks that we considered for both the groups. The *p*-values smaller than 0.01 indicate that our method can identify network differences at a 99% confidence level.

## Results

For each of the 400 subjects, we computed the whole-brain functional connectivity by calculating the Pearson correlation matrix over 1200 time points across 116 anatomical regions resulting in 400 correlation matrices of dimension 116 × 116. Therefore, using our notations, we have *p* = 116 nodes, *q* = *p*(*p* – 1)/2 = 6670 edges, *m*_0_ = *p* – 1 = 115, and *m*_1_ = (*p* – 1)(*p* – 2)/2 = 6555.

### Two-sample test using ETL statistic

Given the 400 correlation matrices of 168 males and 232 females, we aim to check whether the proposed ETL statistic can determine the difference between the groups of males and females. We assume that the male and female edge weights are coming from distributions with cdfs *F_U_* and *F_V_*, respectively. However, these distribution functions are unknown. Therefore, we need to estimate them because the ETL statistic is constructed using these cdfs. To estimate the cdf, we average the male (female) correlation matrices across 168 subjects (232 subjects) and find the empirical cdf based on the averaged 6670 edge weights. The empirical cdfs of the average edge weights of females (in solid black line) and males (in dashed red line) are presented in the left panel of Fig 8. We observe that the empirical cdf corresponding to female is slightly higher than that of male. This suggests a relatively more number of edge weights with smaller values for the female, and a relatively more number of edge weights with bigger values for the male. In other words, the distribution of the female edge weights is slightly positively skewed than the male edge weights. Fig 9 plots the *β*_0_ and *β*_1_ curves of the average female and male networks (calculated in the standard way) and their corresponding estimated counterparts (computed using the expected birth and death values). We observe that the estimated Betti curves well approximate the original Betti curves.

**Fig. 7.**
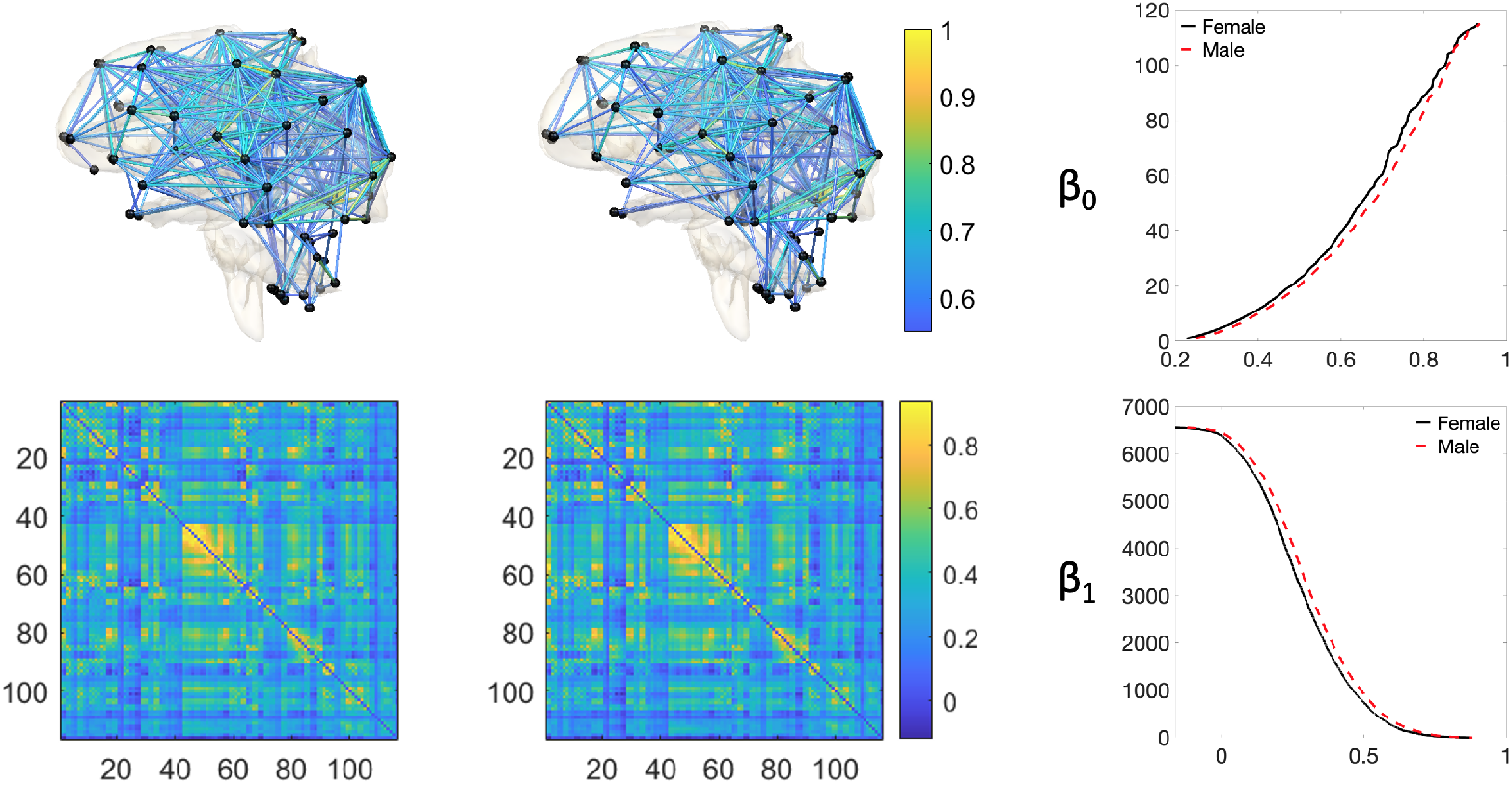
Visualization of the fMRI brain data. Left panel: The average female brain network (top) and the corresponding correlation matrix (bottom). Middle panel: The average male brain network (top) and the corresponding correlation matrix (bottom). Right panel: The *β*_0_ (top) and *β*_1_ (bottom) curves for female (in solid black) and male (in dashed red) brain networks. For a better visualization, we consider a threshold value of 0.5 while plotting the brain networks so that they contain fewer number of edges.

**Fig. 8.**
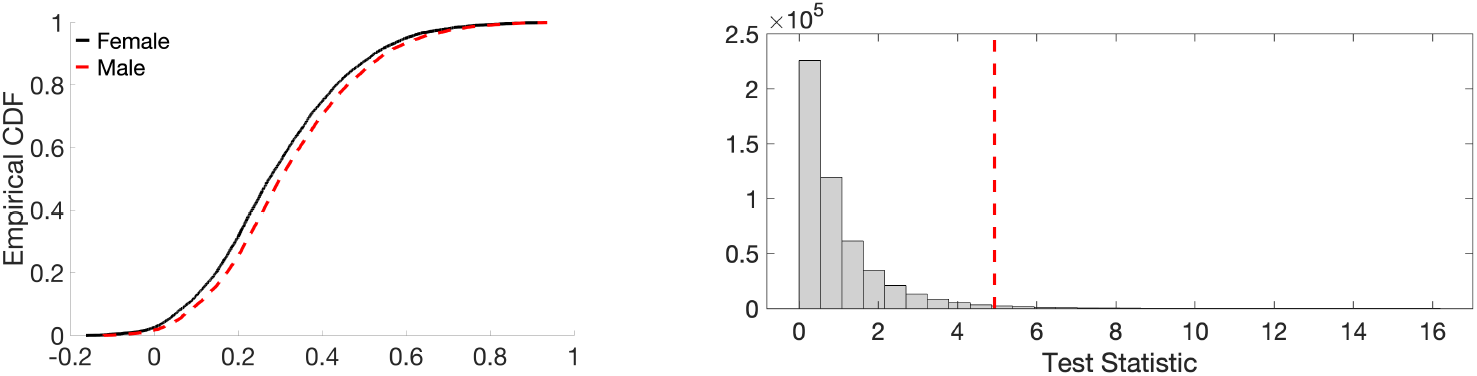
Left: Plot of the empirical cdfs of the average edge weights of females (in solid black line) and males (in dashed red line). Right: Histogram plot of the ETL statistic based on the resting-state fMRI dataset. The dotted red line represents the observed value of the ETL statistic.

**Fig. 9.**
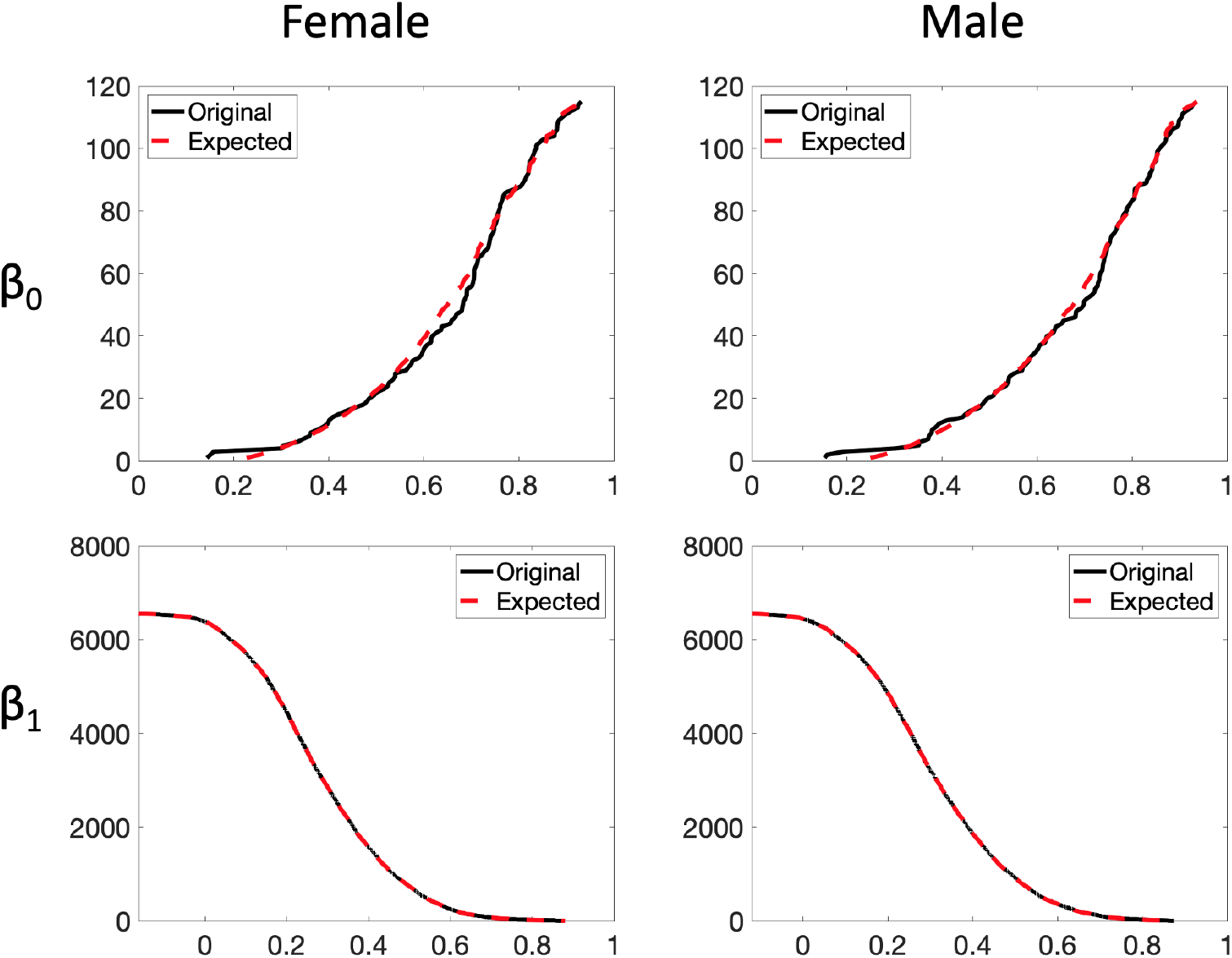
Plots of the original (solid black line) and estimated (dashed red line) *β*_0_ (top) and *β*_1_ (bottom) curves using expected birth and death values for the female (left) and male (right) brain networks.

To conduct the test, we used the permutation test with 500000 random permutations. The observed test statistic is 4.9372 and the *p*-value is 0.0134. The histogram of test statistic is plotted in the right panel of Fig 8. We conclude that, although the weight distributions of the males and females are very close, the proposed ETL statistic can still discriminate them at a 95% confidence level.

### Two-sample test using AUC statistic

We conducted a two-sample test using the method based on the area under *β*_0_ curve. The observed value of the Wilcoxon rank-sum statistic is 48374. The statistic corresponds to the *p*-value of 0.1036. That is, the test fails to discriminate male and female subjects if we use the traditional values of *α*, the level of significance, to be 0.05 or 0.1. However, if we relax this assumption a bit, the test can discriminate males and females at a confidence level of 89.5%.

## Conclusion

The concept of random graphs was first proposed in mid-twentieth century [72] and has been of many researchers’ interest ever since [73–77]. The concepts of TDA tools such as persistent barcodes were extended to handle stochastic cases, which triggered the computation of expected persistent barcodes. However, such computation may require complex theoretical constructs. In this article, we considered a random graph model for which the computation of expected persistent barcodes became simplified by using the *order statistic*.

[32] formulated a topological loss based on the birth and death values of connected components and cycles of a network that provided an optimal matching and alignment at the edge level. In this article, we extended this formulation to a random graph scenario and proposed the expected topological loss (ETL) based on the expected birth and death values. We use the ETL as a test statistic to discriminate between two groups of networks. We validated this method using a simulation study. We showed that the ETL can identify group differences at a 99% confidence level whereas it produces large *p*-values when there is no network differences. We compared the proposed approach with baseline approaches and established an overall superior performance of the proposed method. Further, we considered the area under the Betti curves [68]. This resulted a scalar quantification of the curves which was used to discriminate between the groups. A respective simulation study showed its successful discriminative ability whenever there are network differences. We also applied the developed tools in a resting-state brain fMRI dataset and showed that they can differentiate male and female brain networks.

To calculate the expected persistent barcodes, we computed the unknown distribution using the nonparametric empirical distribution function. However, one may also consider hierarchical or Bayesian parametric models for the edge weights instead. For example, one may consider the edge weights to be drawn from a 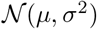 distribution, where the location parameter *μ* and the dispersion parameter *σ*^2^ have a Gaussian and an inverse gamma conjugate prior, respectively. The parameters of the prior distributions will allow flexibility while we can still enjoy the advantages of a parametric model. This direction can be pursued in the future.

## Acknowledgments

We like to thank Shih-Gu Huang of National University of Singapore for providing support for fMRI processing. This study is funded by NIH R01 EB02875 and NSF MDS-2010778.

## References

1. Bassett DS, Sporns O. Network neuroscience. Nature neuroscience. 2017;20(3):353–364.

2. Sporns O. Graph theory methods: applications in brain networks. Dialogues in Clinical Neuroscience. 2018;20(2):111.

3. Bullmore E, Sporns O. Complex brain networks: graph theoretical analysis of structural and functional systems. Nature reviews neuroscience. 2009;10(3):186–198.

4. Stam CJ, Reijneveld JC. Graph theoretical analysis of complex networks in the brain. Nonlinear biomedical physics. 2007;1(1):1–19.

5. Rubinov M, Sporns O. Complex network measures of brain connectivity: uses and interpretations. Neuroimage. 2010;52(3):1059–1069.

6. Chung MK, Lee H, DiChristofano A, Ombao H, Solo V. Exact topological inference of the resting-state brain networks in twins. Network Neuroscience. 2019;3:674–694.

7. Bassett DS. Small-World Brain Networks. The Neuroscientist. 2006;12:512–523.

8. Bassett DS, Porter MA, Wymbs NF, Grafton ST, Carlson JM, Mucha PJ. Robust detection of dynamic community structure in networks. Chaos: An Interdisciplinary Journal of Nonlinear Science. 2013;23(1):013142.

9. Van Den Heuvel MP, Sporns O. Rich-club organization of the human connectome. Journal of Neuroscience. 2011;31(44):15775–15786.

10. Chung MK, Hanson JL, Lee H, Adluru N, Alexander AL, Davidson RJ, et al. Persistent Homological Sparse Network Approach to Detecting White Matter Abnormality in Maltreated Children: MRI and DTI Multimodal Study. MICCAI, Lecture Notes in Computer Science (LNCS). 2013;8149:300–307.

11. Lee H, Kang H, Chung MK, Kim BN, Lee DS. Persistent brain network homology from the perspective of dendrogram. IEEE Transactions on Medical Imaging. 2012;31:2267–2277.

12. Zomorodian AJ, Carlsson G. Computing Persistent Homology. Discrete and Computational Geometry. 2005;33:249–274.

13. Singh G, Memoli F, Ishkhanov T, Sapiro G, Carlsson G, Ringach DL. Topological analysis of population activity in visual cortex. Journal of Vision. 2008;8:1–18.

14. Ghrist R. Barcodes: The persistent topology of data. Bulletin of the American Mathematical Society. 2008;45:61–75.

15. Carlsson G, Memoli F. Persistent clustering and a theorem of J. Kleinberg. arXiv preprint arXiv:08082241. 2008;.

16. Edelsbrunner H, Harer J. Persistent Homology - a Survey. Contemporary Mathematics. 2008;453:257–282.

17. Petri G, Expert P, Turkheimer F, Carhart-Harris R, Nutt D, Hellyer PJ, et al. Homological scaffolds of brain functional networks. Journal of The Royal Society Interface. 2014;11:20140873.

18. Sizemore AE, Giusti C, Kahn A, Vettel JM, Betzel RF, Bassett DS. Cliques and cavities in the human connectome. Journal of computational neuroscience. 2018;44:115–145.

19. Lee H, Chung MK, Kang H, Choi H, Kim YK, Lee DS. Abnormal hole detection in brain connectivity by kernel density of persistence diagram and Hodge Laplacian. In: IEEE International Symposium on Biomedical Imaging (ISBI); 2018. p. 20–23.

20. Bubenik P. The persistence landscape and some of its properties. In: Topological Data Analysis. Springer; 2020. p. 97–117.

21. Chung MK, Lee H, Solo V, Davidson RJ, Pollak SD. Topological distances between brain networks. International Workshop on Connectomics in Neuroimaging. 2017; p. 161–170.

22. Lee H, Chung MK, Kang H, Kim BN, Lee DS. Computing the shape of brain networks using graph filtration and Gromov-Hausdorff metric. MICCAI, Lecture Notes in Computer Science. 2011;6892:302–309.

23. Songdechakraiwut T, Shen L, Chung MK. Topological learning and its application to multimodal brain network integration. Medical Image Computing and Computer Assisted Intervention (MICCAI). 2021; p. in press, http://pages.stat.wisc.edu/~mchung/papers/song.2021.MICCAI.pdf.

24. Anand DV, Chung MK. Hodge-Laplacian of Brain Networks and Its Application to Modeling Cycles. arXiv preprint arXiv:211014599. 2021;.

25. Berry E, Chen YC, Cisewski-Kehe J, Fasy BT. Functional summaries of persistence diagrams. Journal of Applied and Computational Topology. 2020;4(2):211–262.

26. Chazal F, Fasy BT, Lecci F, Rinaldo A, Wasserman L. Stochastic convergence of persistence landscapes and silhouettes. In: Proceedings of the thirtieth annual symposium on Computational geometry; 2014. p. 474–483.

27. Bubenik P. Statistical topological data analysis using persistence landscapes. Journal of Machine Learning Research. 2015;16:77–102.

28. Biscio CA, Møller J. The accumulated persistence function, a new useful functional summary statistic for topological data analysis, with a view to brain artery trees and spatial point process applications. Journal of Computational and Graphical Statistics. 2019;28:671–681.

29. Chen YC, Wang D, Rinaldo A, Wasserman L. Statistical analysis of persistence intensity functions. arXiv preprint arXiv:151002502. 2015;.

30. King JB, Prigge MB, King CK, Morgan J, Weathersby F, Fox JC, et al. Generalizability and reproducibility of functional connectivity in autism. Molecular Autism. 2019;10(1):1–23.

31. Blinowska KJ, Kaminski M. Functional brain networks: random, “small world” or deterministic? PloS one. 2013;8(10):e78763.

32. Songdechakraiwut T, Chung MK. Topological learning for brain networks. 2020; p. arXiv:2012.00675.

33. Bollobás B, Béla B. Random graphs. 73. Cambridge university press; 2001.

34. Frieze A, Karoński M. Introduction to random graphs. Cambridge University Press; 2016.

35. Gayet D, Welschinger JY. Lower estimates for the expected Betti numbers of random real hypersurfaces. Journal of the London Mathematical Society. 2014;90(1):105–120.

36. Salepci N, Welschinger JY. Tilings, packings and expected Betti numbers in simplicial complexes. arXiv preprint arXiv:180605084. 2018;.

37. Wigman I. On the expected Betti numbers of the nodal set of random fields. Analysis & PDE. 2021;14(6):1797–1816.

38. Wilks SS. Order statistics. Bulletin of the American Mathematical Society. 1948;54(1):6–50.

39. Rényi A. On the theory of order statistics. Acta Mathematica Academiae Scientiarum Hungarica. 1953;4(3-4):191–231.

40. David HA, Nagaraja HN. Order statistics. John Wiley & Sons; 2004.

41. Arnold BC, Balakrishnan N, Nagaraja HN. A first course in order statistics. SIAM; 2008.

42. Ahsanullah M, Nevzorov VB, Shakil M. An introduction to order statistics. vol. 8. Springer; 2013.

43. Balakrishnan N, Cohen AC. Order statistics & inference: estimation methods. Elsevier; 2014.

44. Van Essen DC, Ugurbil K, Auerbach E, Barch D, Behrens TEJ, Bucholz R, et al. The Human Connectome Project: a data acquisition perspective. NeuroImage. 2012;62:2222–2231.

45. Glasser MF, Sotiropoulos SN, Wilson JA, Coalson TS, Fischl B, Andersson JL, et al. The minimal preprocessing pipelines for the Human Connectome Project. Neuroimage. 2013;80:105–124.

46. Bzdok D, Varoquaux G, Grisel O, Eickenberg M, Poupon C, Thirion B. Formal models of the network co-occurrence underlying mental operations. PLoS computational biology. 2016;12:e1004994.

47. Desikan RS, Ségonne F, Fischl B, Quinn BT, Dickerson BC, Blacker D, et al. An automated labeling system for subdividing the human cerebral cortex on MRI scans into gyral based regions of interest. NeuroImage. 2006;31:968–980.

48. Hagmann P, Kurant M, Gigandet X, Thiran P, Wedeen VJ, Meuli R, et al. Mapping human whole-brain structural networks with diffusion MRI. PLoS One. 2007;2(7):e597.

49. Arslan S, Ktena SI, Makropoulos A, Robinson EC, Rueckert D, Parisot S. Human brain mapping: A systematic comparison of parcellation methods for the human cerebral cortex. NeuroImage. 2018;170:5–30.

50. Tzourio-Mazoyer N, Landeau B, Papathanassiou D, Crivello F, Etard O, Delcroix N, et al. Automated anatomical labeling of activations in SPM using a macroscopic anatomical parcellation of the MNI MRI single-subject brain. NeuroImage. 2002;15:273–289.

51. Power JD, Barnes KA, Snyder AZ, Schlaggar BL, Petersen SE. Spurious but systematic correlations in functional connectivity MRI networks arise from subject motion. NeuroImage. 2012;59:2142–2154.

52. Van Dijk KRA, Sabuncu MR, Buckner RL. The influence of head motion on intrinsic functional connectivity MRI. NeuroImage. 2012;59:431–438.

53. Satterthwaite TD, Wolf DH, Loughead J, Ruparel K, Elliott MA, Hakonarson H, et al. Impact of in-scanner head motion on multiple measures of functional connectivity: relevance for studies of neurodevelopment in youth. NeuroImage. 2012;60:623–632.

54. Caballero-Gaudes C, Reynolds RC. Methods for cleaning the BOLD fMRI signal. NeuroImage. 2017;154:128–149.

55. Edelsbrunner H, Harer J. Computational topology: An introduction. American Mathematical Society; 2010.

56. Mileyko Y, Mukherjee S, Harer J. Probability measures on the space of persistence diagrams. Inverse Problems. 2011;27(12):124007.

57. Vallender S. Calculation of the Wasserstein distance between probability distributions on the line. Theory of Probability & Its Applications. 1974;18(4):784–786.

58. Mi L, Zhang W, Gu X, Wang Y. Variational Wasserstein clustering. In: Proceedings of the European Conference on Computer Vision (ECCV); 2018. p. 322–337.

59. Mi L, Zhang W, Wang Y. Regularized Wasserstein means for aligning distributional data. In: Proceedings of the AAAI Conference on Artificial Intelligence. vol. 34; 2020. p. 5166–5173.

60. Panaretos VM, Zemel Y. Statistical aspects of Wasserstein distances. Annual review of statistics and its application. 2019;6:405–431.

61. Kolouri S, Zou Y, Rohde GK. Sliced Wasserstein kernels for probability distributions. In: Proceedings of the IEEE Conference on Computer Vision and Pattern Recognition; 2016. p. 5258–5267.

62. Mosteller F. On some useful “inefficient” statistics. In: Selected Papers of Frederick Mosteller. Springer; 2006. p. 69–100.

63. Duong T, Hazelton ML. Cross-validation bandwidth matrices for multivariate kernel density estimation. Scandinavian Journal of Statistics. 2005;32(3):485–506.

64. Thompson PM, Cannon TD, Narr KL, van Erp T, Poutanen VP, Huttunen M, et al. Genetic influences on brain structure. Nature Neuroscience. 2001;4:1253–1258.

65. Zalesky A, Fornito A, Harding IH, Cocchi L, Yücel M, Pantelis C, et al. Whole-brain anatomical networks: Does the choice of nodes matter? NeuroImage. 2010;50:970–983.

66. Nichols TE, Holmes AP. Nonparametric permutation tests for functional neuroimaging: A primer with examples. Human Brain Mapping. 2002;15:1–25.

67. Winkler AM, Ridgway GR, Douaud G, Nichols TE, Smith SM. Faster permutation inference in brain imaging. NeuroImage. 2016;141:502–516.

68. Xu F, Garai S, Chung M, Caciagli L, Saykin AJ, Bassett DS, et al. Identifying topological changes of structural connectome in MCI and AD through persistent homology. In preperation. 2021;.

69. Haynes W. In: Dubitzky W, Wolkenhauer O, Cho KH, Yokota H, editors. Wilcoxon Rank Sum Test. New York, NY: Springer New York; 2013. p. 2354–2355. Available from: https://doi.org/10.1007/978-1-4419-9863-7_1185.

70. Cohen-Steiner D, Edelsbrunner H, Harer J. Stability of Persistence Diagrams. Discrete and Computational Geometry. 2007;37:103–120.

71. Chazal F, Cohen-Steiner D, Guibas LJ, Mémoli F, Oudot SY. Gromov-Hausdorff Stable Signatures for Shapes using Persistence. In: Computer Graphics Forum. vol. 28; 2009. p. 1393–1403.

72. Erdös P, Rényi A. On the evolution of random graphs. Bull Inst Internat Statist. 1961;38:343–347.

73. Kovalenko I. Theory of random graphs. Cybernetics. 1971;7(4):575–579.

74. Barabási AL, Albert R. Emergence of scaling in random networks. science. 1999;286(5439):509–512.

75. Karoński M, Scheinerman ER, Singer-Cohen KB. On random intersection graphs: The subgraph problem. Combinatorics, Probability and Computing. 1999;8(1-2):131–159.

76. Chung F, Lu L. Connected components in random graphs with given expected degree sequences. Annals of combinatorics. 2002;6(2):125–145.

77. Leskovec J, Chakrabarti D, Kleinberg J, Faloutsos C. Realistic, mathematically tractable graph generation and evolution, using kronecker multiplication. In: European conference on principles of data mining and knowledge discovery. Springer; 2005. p. 133–145.

